# Screening Peptide Drug Candidates to Neutralize Whole Viral Agents : A Case study with SARS-CoV-2 Virus

**DOI:** 10.1101/2023.10.22.563490

**Authors:** Cemile Elif Özçelik, Cemre Zekiye Araz, Özgür Yılmaz, Sevgi Gülyüz, Aykut Özkul, Urartu Özgür Şafak Şeker

## Abstract

Covid19 pandemic revealed the reality for the need of therapeutic and pharmaceutical molecule development in a short time with different approaches. Although the enhancement of immunological memory by vaccination was the quicker and robust strategy, still medication is required for immediate treatment for a patient. For this purpose, one of the approaches is developing new therapeutic molecule development like peptide-based drugs. Also, peptides can be used developing other molecules like nanobodies. Here, M13 phage display library was used for selecting SARS-CoV-2 interacting peptides for developing a neutralizing molecule for further use. Biopanning was applied with four iterative cycles to select phages displaying different 12-amino acid-long peptides. Then, the M13 phage genomic region where peptide sequences expressed were analyzed and sequences were obtained. Randomly selected peptide sequences were synthesized by solid-state peptide synthesis method. These peptides were analyzed by quartz crystal microbalance method in terms or peptide interaction capacity with specifically wild-type S protein. Next, QCM data was further validated by enzyme-linked immunosorbent assay (ELISA) in order to check peptides according to their neutralizing capacity rather than binding to S1 protein. The results showed that, phage display served an opportunity for selecting peptides which can be used and developed further as pharmaceutical molecules. More specifically, scpep3, scpep8 and scpep10 had both binding and neutralizing capacity for S1 protein as a candidate for therapeutic molecule.

---

The severe acute respiratory syndrome coronavirus 2 (SARS-CoV-2) gave rise to a worldwide pandemic crisis and played havoc with public health by causing coronavirus disease 2019 (Covid19)^1^. The potential of the spread of the disease easily, that affects the daily life, changed the perspectives for developing fast and effective treatment approaches and unearthed the possible life threating disease spreads^2^.

As of now, both Covid19 pandemic and the possible future outbreaks catch the eye for developing new therapeutic and detection approaches and molecules^3^. Since only vaccination is the best option to develop immunity to SARS-CoV-2 infection, the protection of vaccination against different variants is not only way to eliminate such a pandemic^4^. There is always a need for discovering and developing therapeutic agents as medications for patients during disease to relieve from the symptoms and treat the disease^5^. With this purpose in mind, Covid19 drug molecule developments gathered around two distinct targets, which are host proteins associated with viral particle interaction and viral proteins^6–8^. Viral inhibitor developments targeting host proteins are focused mainly on angiotensin-converting enzyme 2 (ACE2), and cellular proteases, which are furin, transmembrane protease serine2, ADAM10, ADAM17, and cathepsin, as well as several other host proteins like caspase-6, CD147, farnesoid X receptor, and eukaryotic translation elongation factor 1A^9–19^. The main approach for antiviral agent development is inhibiting the entry of SARS-CoV-2 into host cells such as through viral protease inhibition^20^. Antiviral agent development centered on the viral proteins intends to inhibit some of the events in the SARS-CoV-2 life cycle^21^. Spike (S) protein inhibition, RNA synthesis inhibition, viral particle assembly blockage, and hindering proteolytic processes are the main focus of blocking viral particles during infection^22–28^. Neutralizing the viral particles during infection is highly effective for inhibition of SARS-CoV-2 and preventing disease progression^29^. More than 700 agents have been registered as antiviral molecules against SARS-CoV-2. The majority of the antiviral molecules are small molecules (53%). Next in line is antibodies (33%), followed by peptide inhibitors (4%) and the others. Although the number of antiviral agents under preclinical or clinical trials was satisfactory, about 90% of the molecules failed to further stages for approval. This situation proves the need for developing more molecules against SARS-CoV-2^30^.

Commencing with this juncture, the rapid drug molecule discovery or neutralizing agent development can be achieved by phage display library. In this regard, we aimed to select interacting peptides against inactivated SARS-CoV-2 particles. For selecting candidate peptides, the library used in this study expressed 12 amino-acid long peptides at M13KE pIII coat protein. To eliminate selection of plastic-binding peptides in panning method, preselection, which has the same steps as panning method except library elements were collected from the unbound phages, was applied (Figure 1A). To achieve more specific phage particles for inactivated viral particles, unbound phages were eluted and amplified. Thereby, SARS-CoV-2 specific library was obtained by preselection method application. Further, the selection of interacting peptides was achieved by applying panning method after four iterative cycles.

**Figure 1:**
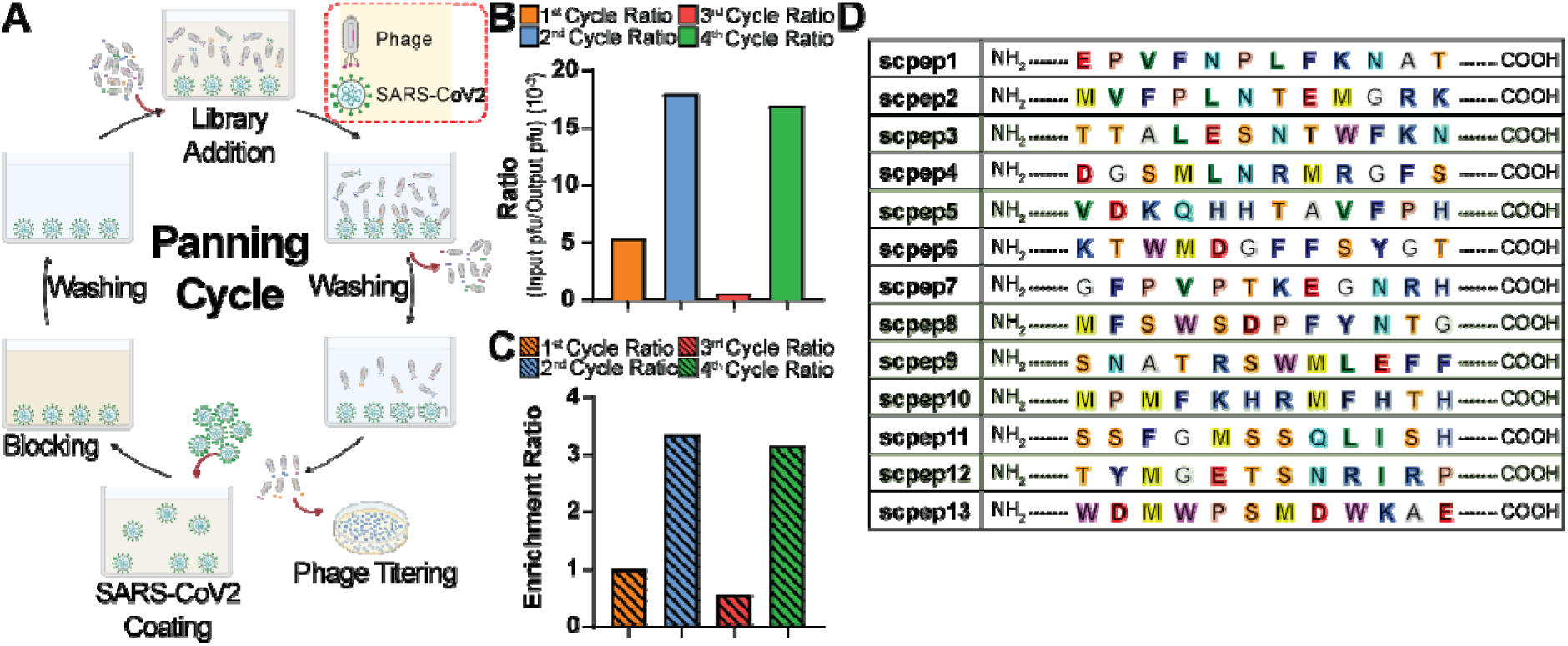
Using phage display library provided an option to select peptides which were highly specific to inactivated SARS-CoV-2. (A) The one cycle of panning was achieved by immobilization of inactivated SARS-CoV-2 which was followed by blocking, phage library addition and bound-phage elution. After blocking and incubation with phage library, washing steps were applied. At the end of the cycle, phage titering and phage amplification were done in order to calculate the eluted phage amount and for next panning cycles with reduced number of phage clones, respectively. The figure created with BioRender.com. (B) Following the completing one panning cycle, phage particle ratio was calculated. (C) Each cycle provided an enrichment ratio which was important to specify the more interacted phage clone expansion. Although enrichment ratio of the third cycle possessed the lowest enrichment ratio, the overall phage particle specificity was increased by removal of unbound or mildly bound-phage particles according to the result of the previous cycle except the third cycle. (D) When four iterative panning cycles were completed, the genomic region where peptide sequences expressed were analyzed and several candidate peptide sequences were determined.

As a starting, 10^9^ phage particles and fixed volume of viral particles in the same sample were used. After the first round of selection finished, the eluted phages were amplified to produce a phage library for the second round of panning. This process was continued for four panning cycles in total. Also, after each cycle, the number of phage particles in phage elution and amplified phage samples were determined by tittering. According to the determined particle amount, the ratio was calculated by dividing phage particle amount in elution as output phage amount by the library size (pfu/mL) used each panning cycle as output phage amount, which was 10^9^. Based on the data procured by phage tittering, there was an increase in the ratio derived from tittering of eluted phages and amplified phages from the elution between the first and the fourth rounds of panning (Figure 1B). Specifically, as expected, the second round of panning ratio result was higher than the first round of panning ratio. However, there was a decrease in the ratio after the third round of panning. We suspect the non-specifically bound phages were washed out mostly at this step. Still, at the end of the fourth round, there was an increase in ratio depending on the third round. When the ratio results were compared, there was a 3.35-fold increase in the ratio between the first and the second rounds and a 5.7-fold increase between the third and fourth rounds, which was satisfactory to continue the selection of phage-forming units from the last elution tittering.

Likewise, enrichment ratios were calculated by dividing the ratio of the round by the ratio result of the first-round panning. The enrichment ratio results were similar to the ratio results (Figure 1C). As expected, there were raises in the enrichment ratios in the second and fourth rounds of panning. As in ratio data, it was foreseen that the decrease in enrichment ratio was observed in the result of the third round after the second round. Nevertheless, the final result showed that the final phage elution was enriched in terms of more interacting phage particle existence after completing 4 cycles of the panning protocol.

Once panning cycles were completed, eluted phage tittering products were used to select single phage colonies for sequencing. As M13KE bacteriophage pIII coat protein was used to display the library, *pIII* gene regions of the selected phage colonies were analyzed by Sanger sequencing method (Genewiz). As specified by the Sanger sequencing data, thirteen peptide sequences were determined (Figure 1D). Before synthesizing the peptides, all were analyzed by using The Coronavirus Antibody Database to determine the similarities with the third complementary determining region (CDR3) of the antibody heavy chain^31^. This region is vital for specific antibody binding^32^. Also, peptides were checked for probability for polystyrene binding capacity by using SAROTUP: Target-Unrelated Peptides Scanners^33,34^. As per analyses, the amount of 12 amino acid-long CDRs as well as similarities in the database were obtained. Furthermore, the charge at pH 7.0 was calculated. Likewise, each peptide was checked for the probability for polystyrene binding. Even though the polystyrene binding probabilities were obtained, the peptides were selected randomly regardless from such information. The reason of random selection of peptides for further analyses was to evaluate the importance of such information as well as the power of pre-selection should eliminate the polystyrene bound-phage particles. This provided an option for checking peptides for interaction with a target without selecting them according to probability to polystyrene binding. Besides, our random selection basically was focused on the randomness of other information like sequence the 12-amino acid long CDRs in the databank, identity and pH.

When the peptide synthesis was completed by solid-state synthesis, the interaction of peptides with S protein as well as the neutralization activity of peptides for S protein interaction with Ace2-Fc protein, first S protein and Ace2-Fc protein were expressed in Expi298 and purified by HisTag purification by using 5mL Cytiva HisTrap™ Excel column.

After synthesizing peptides, the peptide interaction studies with S protein, as well as the neutralization activity of peptides were done by HisTagged pure S protein and Ace2-Fc protein. To achieve the expression and purification, S protein and Ace2-Fc protein genes were cloned into Ap2u2-mcherry plasmid, a gift from Christien Merrifield (Addgene plasmid # 27672; http://n2t.net/addgene:27672; RRID:Addgene_27672)^35^. Subsequent to expression and purification of S protein and Ace2-Fc, the first characterization studies depended on the interaction of peptides with S proteins. For this purpose, quartz crystal microbalance (QCM) analysis was done for each synthesized peptide. This analysis demonstrates the interaction of two molecules by immobilizing one molecule on a gold chip and delivering the second molecule onto the chip surface by a peristaltic pumping system.

During the analysis, the surface of the gold chip was coated with 11-Mercaptoundecanoic acid (11-MUA) and activated by EDC/NHS coupling reaction. The activated surface was then coated by S protein and deactivated by Ethanolamine HCl. For each type of molecule or chemical addition to the QCM chamber, the sensor recorded ΔF (Hz) changes. EDC, S protein, and ethanolamine HCL addition decreases the ΔF values, although NHS addition increases the ΔF values. The chamber was washed with 1X PBS between each buffer or molecule addition to remove excessive molecules on the gold chip. After chip preparation was achieved by S protein coating, peptides were delivered to the chip surface several times in one measurement. Subsequent to each peptide addition to the system for one measurement, the chip was washed with 1X PBS. The decrease in the value of ΔF meant there was a mass accumulation onto the chip, and an increase in the value of ΔF represents the removal of the unbound or excessive peptide molecules.

Depending on this information, S protein coating was successfully achieved by obtaining the desired decrease or increase in ΔF values. In the case of scpep3, the accumulated S protein mass on the chip was 62.48 ng/cm^2^. Following S protein immobilization, scpep3 was introduced to QCM chamber repetitively. At the beginning of scpep3 reaching to the chip surface, there was an increase in ΔF, which was unexpected. After several scpep3 addition, the ΔF change gave proper results with 6.62 ng/cm^2^ mass accumulation according to starting and final ΔF5 results.

For scpep5, the gold chip was prepared by accumulation 629.05 ng/cm^2^ S protein on the chip. scpep5 addition, on the other hand, accumulated on the gold chip at the very first addition to the chamber. However, washing steps increases the ΔF up to positive values. Each scpep5 addition decreases the ΔF, although frequency values stand in positive values. Thus, these results fell short of expectations by giving negative measurements for mass accumulation.

In the case of scpep8 interaction evaluation, S protein coating succeeded with 820.77 ng/cm^2^ of mass accumulation. This was followed by scpep8 addition. For the first addition of scpep8 to the QCM chamber, ΔF got more negative values. After washing step, some of the unbound scpep8 molecules seem removed on the chip surface. Repetition of the addition of scpep8, followed by washing steps, increased ΔF values slightly. Still, the overall change in frequency values stayed in negative values with 7.81 ng/cm^2^ of mass accumulation, which was satisfactory.

Regarding with scpep9 QCM analysis, S protein coating was done by accumulating 89.28 ng/cm^2^ protein mass onto the gold chip. scpep9 delivery to S protein coated chip gave unusual ΔF profiles by increasing the frequency change values and then decreasing ΔF profile was observed. Because of the exclusive ΔF profile, there was only 14.04 ng/cm^2^ of mass accumulation was calculated by considering only from the point where ΔF5 became zero second time then started to give negative values and final ΔF5 values.

When it comes to the interaction of scpep10 with S protein, first 274.65 ng/cm^2^ S protein coating was accomplished onto gold chip. Then, scpep10 was added to the tubing system of QCM chamber for measuring ΔF. The ΔF values became more negative during scpep10 addition. After the washing step, the ΔF values were obtained in a more proper profile with 10.73 ng/cm^2,^ peptide accumulation, which was actually expected for all interacted peptide molecules against an immobilized target. This result was highly promising and satisfactory for further evaluation.

Finally, scpep12 was checked for interaction with S protein. To do so, 506.43 ng/cm^2^ S protein was immobilized on gold chip. As with the results of scpep5 QCM data, scpep12 also gave an overall ΔF increase, although each scpep12 addition has resulted in a decline in ΔF profile. The positive ΔF values did not give a mass accumulation.

**Figure 2:**
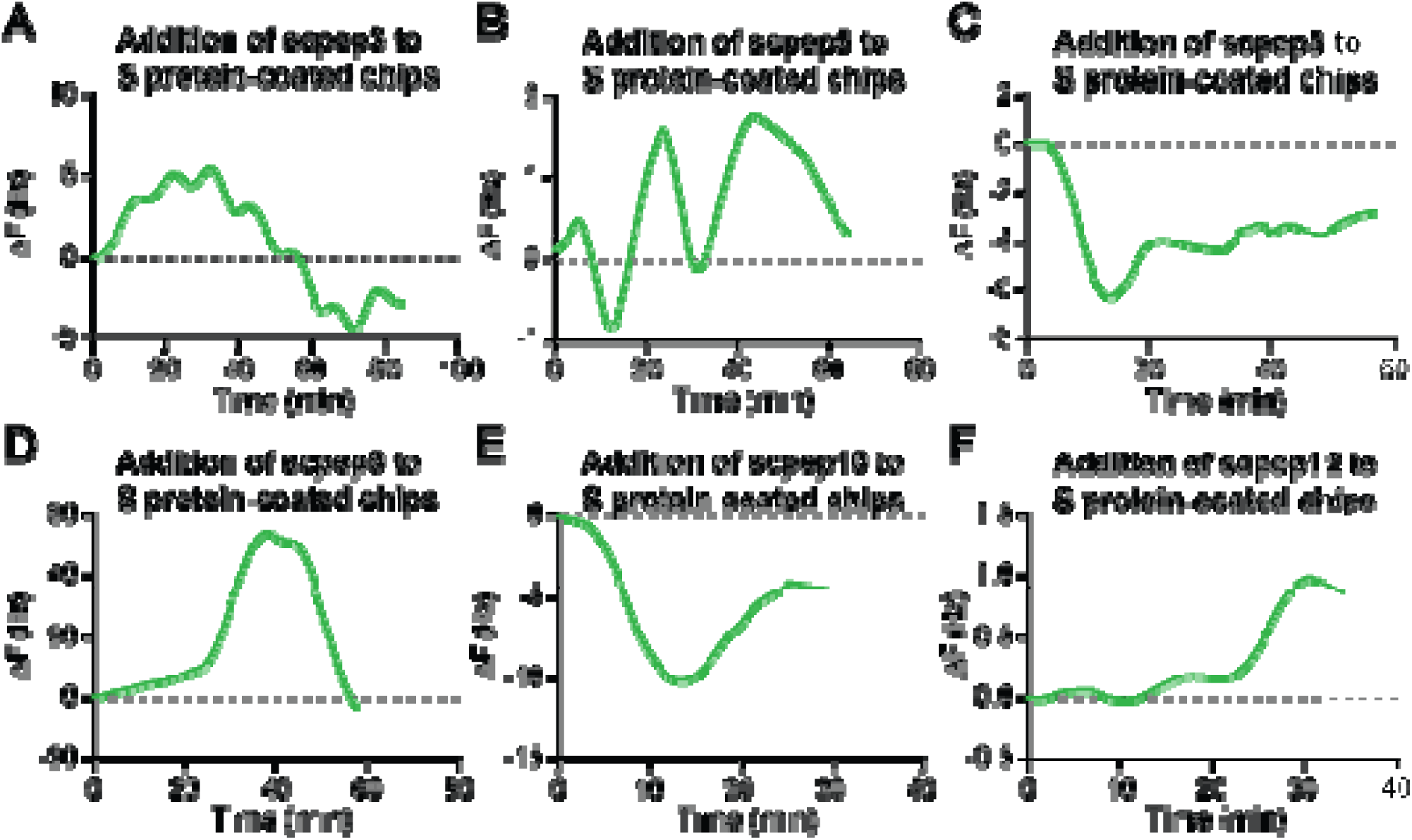
Peptide-S protein interactions were determined by QCM analysis according to change in the frequencies. (A) scpep3 created unexpected ΔF profile at the beginning although it continued with a proper decrease in ΔF by increasing peptide accumulation onto S protein-coated chip. (B) scpep5 addition demonstrated decrease in ΔF. However, each washing step removed a mass from the chip. (C) scpep8 provided more optimal ΔF as peptide accumulation occurred for each addition. (D) scpep9 addition showed a deposition profile in ΔF. Also, washing step created an unexpected decrease in ΔF. (E) scpep10 exhibited the best ΔF profile by a decline in the ΔF plot during peptide addition and slightly incline of the line representing ΔF. (F) scpep12 addition created small oscillations in ΔF at the beginning which was continued with a deposition profile for ΔF.

According to QCM results, scpep8 and scpep10 gave encouraging outcomes for further analysis. Besides, scpep3 catch the appropriate frequency change profile even though the early records have demonstrated strange increase in frequency change. Also, the unexpected and unusual results observed for other peptides required another analysis. After detecting the binding and the interaction between peptides and immobilized S protein, the peptides were analyzed for neutralization activities in order to determine which peptide could be used as development of therapeutic purposes or other applications like detection. To achieve this, enzyme-linked immunosorbent assay (ELISA) was preferred to evaluate neutralization activities of peptides.

In ELISA, the main aim was to detect the neutralization activity of peptides upon constant (Fc) domain of immunoglobulin G (IgG)-tagged human Ace2 protein by blocking the interaction of S1 protein with Ace2-Fc. Ace2-Fc-coated plates were prepared for each peptide as triplicates. The expectation was that the color should be changed to yellow, and the OD_450_ value was maximized if there was no or low-level neutralization activity of peptides for horseradish peroxidase (HRP)-conjugated S1 protein. Since the absence of neutralization activity of peptides could not disrupt the interaction of S1-HRP with Ace2-Fc protein, the HRP substrate would change the color and increase absorbance. If there was any interaction of peptides with HRP-conjugated S1 protein, then the expectation was that peptides were supposed to block S1 protein binding capacity for Ace2-Fc. Then, peptide-S1-HRP conjugates could not interact with Ace2-Fc, and there would be no or low level of HRP activity after the substrate addition since peptide-S1 complexes were removed and washed away before substrate addition (Supplementary Figure S3).

As a negative control, Ace2-Fc coated plaques were checked with only HRP-conjugated S1 protein, which was supposed to bind to Ace2-Fc and to give a maximum OD_450_ value. For positive controls, S1 protein was incubated with anti-S RBD antibody and added to Ace2-Fc coated plaques. Thus, blocked RBD region of S1 protein could not interact with Ace2-Fc protein because of anti-S RBD antibody. Based on this workflow and experimental design, each peptide was evaluated for peptide-S1 interaction in terms of low OD_450_ value and increased neutralization activity by using S1-HRP conjugates from wild-type SARS-CoV-2, B.1.617.2 and B.1.1.7 variants (Figure 3).

**Figure 3:**
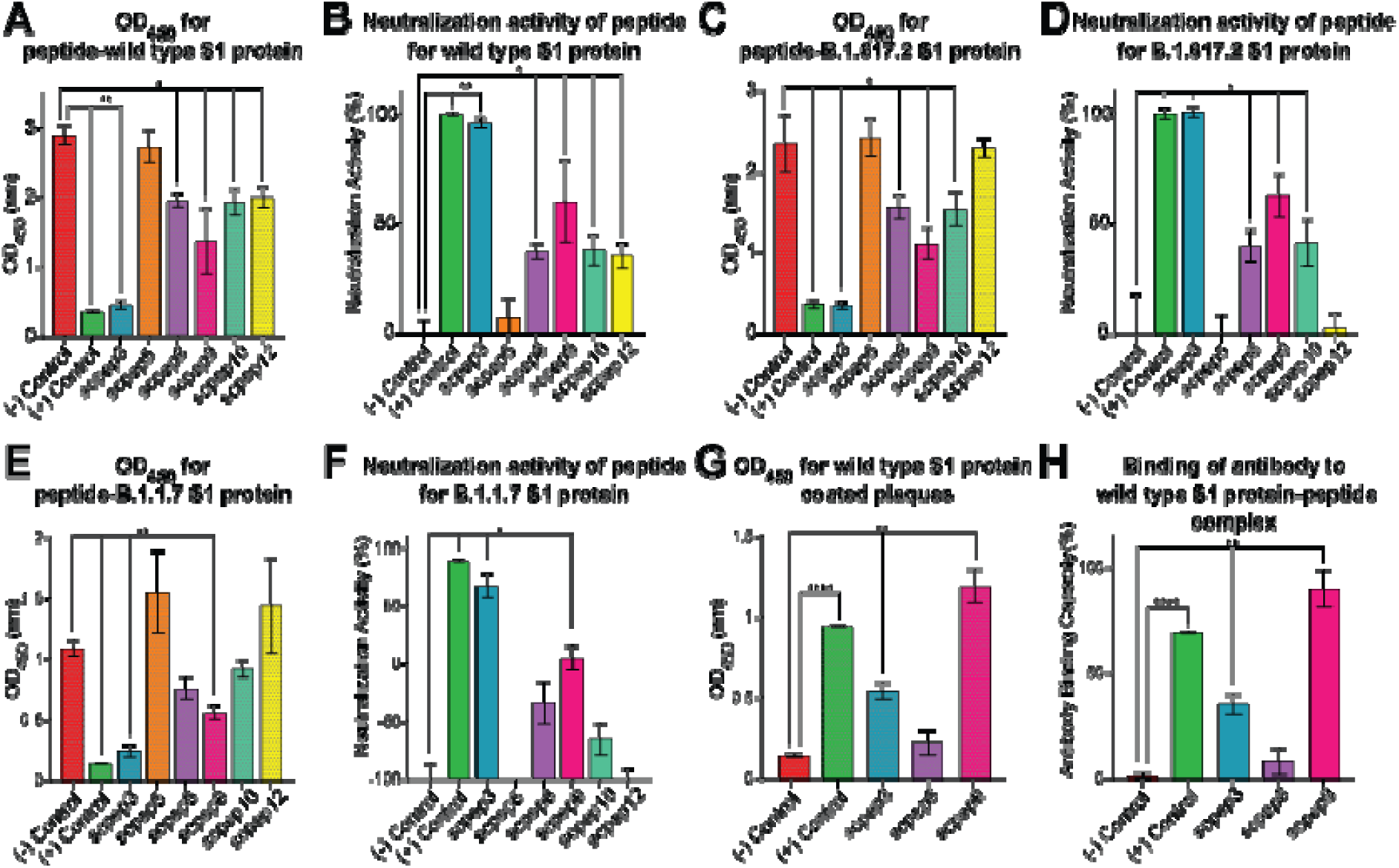
ELISA was applied for all peptides to support the binding capabilities of the peptides with a neutralization activity. ELISA was preferred whether peptides could be used as neutralizing therapeutics or not. (A-B) Peptides were analyzed against Ace2-Fc by mixing with HRP-conjugated S1 protein of wild-type SARS-CoV-2. For each sample, absorbance values at 450 nm were measured. According to this, neutralization activities were calculated by normalizing the data. (C-D) By using B.1.617.2 S1 proteins were conjugated with HRP and mixed with each peptide samples in order to detect OD_450_ values which were derived to neutralization activities. (E-F) S1 protein of B1.1.2 variant was used to characterize the neutralization activities of the peptides by measuring the OD_450_ values which were then converted to demonstrate neutralization activities. (G-H) For evaluating the peptides whether affecting the anti-S RBD antibody interaction with wild-type S1 protein, OD_450_ values for scpep3, scpep8, and scpep9 were measured. Then, antibody binding capacities were determined in the existence of these three peptides. All of the ELISA measurements were undergone T-test for determining significant differences from negative controls. The significances were determined by GP: 0.1234 (ns), 0.0332 (*). 0.0021 (**), 0,0002 (***), 0.0001 (****).

In the case of using wild-type S1 protein, scpep3 gave the lowest OD_450_ value as positive control which were statistically significant from negative control (Figure 3A). On the other hand, scpep5 possessed close OD_450_ value to negative control with no significance from negative control. For scpep8, scpep9, scpep10 and scpep12, OD_450_ values possessed acceptable significant difference from the OD_450_ value of negative control. As expected, scpep3 showed high % of neutralization activity than the others (Figure 3B). scpep5 showed no specific neutralization activity. The average neutralization activities were obtained with scpep8, scpep9, scpep10 and scpep12. With the comparison between QCM analysis data and ELISA data, scpep3, scpep8 and scpep10 were expected to show neutralization activity for wild-type S1 protein. Also, as the peptides selected against inactive wild-type SARS-CoV-2 particles, neutralization activity for wild-type S1 protein was expected with a great possibility, which was proven with ELISA data.

Other than wild-type S1 protein, the neutralization activities of peptides for S1 proteins of B.1.617.2 and B.1.1.7 were evaluated with the intent of observing neutralization activities of peptides the other variants. ELISA OD_450_ measurements were obtained by using S1 protein of B.1.617.2, which were similar with the results obtained from ELISA done by wild-type S1 protein (Figure 3C). More specifically, scpep3 showed neutralization activity like positive control. The only difference between results obtained from ELISA achieved by wild-type and B.1.617.2 variant was that the significance from negative control were slightly lower in OD_450_ values for B.1.617.2 S1 protein. As like the results of scpep5 for wild-type S1 protein, OD_450_ value for scpep5 higher than the others and highly close to negative control OD_450_ value. Beside scpep5, scpep12 also revealed a moderately high OD_450_ value with no significant difference from the negative control.

For the neutralization activity of the peptides, again, scpep3 showed great neutralization for B.1.617.2 S1 protein against Ace2-Fc (Figure 3D). Likewise, scpep8, scpep9, and scpep10 were effective for neutralizing S1 protein. Conversely, scpep3 and scpep12 had no neutralizing capability for S1 protein.

Concerning the result of ELISA using B.1.1.7 S1 protein, the OD_450_ values started to give different results (Figure 3E). Still, scpep3 gave a low OD_450_ value like the result obtained from wild-type S1. In addition to this, scpep9 created a complex with B.1.1.7 S1 protein and produced a slightly low OD_450_ value. However, scpep8 and scpep10 showed no significant difference than the negative control, which was otherwise for the data obtained from wild-type and B.1.617.2 S1 proteins. Similarly, scpep5 and scpep12 possessed a high value for OD_450_. Starting from this point of view, the neutralization activities were also calculated for peptides with B.1.1.7 S1 protein (Figure 3F). As the perspective developed with OD_450_ values, scpep3 had a neutralizing activity for B.1.1.7 S1 protein with an acceptable significant difference from the negative control. Furthermore, scpep9 demonstrated a neutralizing capability for S1 protein. In contrast, especially scpep5 and scpep12, as well as scpep8 and scpep10, also showed no neutralizing function for B.1.1.7 S1 protein.

As an additional verification of the neutralizing activity of the peptides, ELISA was also applied by covering the plates with wild-type S1 protein. In the experimental setup, peptides were added onto S1-coated plate surface, and then anti-S RBD antibody was appended to the wells. To check the anti-S RBD antibody interaction with S1 protein interacted with peptides, mouse anti-human IgG antibody was introduced to the analysis. As a positive control, there was no peptide addition in the workflow. To obtain the negative result, S1 protein-coated wells were only blocked with 3% BSA, and then mouse anti-human IgG antibody was added to the wells. According to this design, positive control would give high OD_450_ results as well as high neutralizing activity (Figure 3G). Similarly, negative controls were expected to give OD_450_ value as small as possible. For this experiment, scpep3, scpep8, and scpep9 were selected to use since scpep3 gave high neutralization activity for all S1 samples, and scpep8 gave moderate neutralizing capacity. Also, scpep9 possessed a neutralization effect for wild-type and B1.617.2 S1 protein, although there was no neutralization activity for B.1.1.2 S1 protein. By following this information, ELISA gave significantly high OD_450_ results from negative control. In the case of interaction of peptides and blocking of the recognition with anti-S RBD antibody, scpep9 gave highest OD_450_ value, meaning that scpep9 enhances the recognition of S1 protein by anti-S RBD antibody (Figure 3G-H). For scpep3, there was a mild increase in OD_450_ value. scpep8 addition has caused an insignificant result for OD_450_. According to this, scpep8 eliminated the anti-S RBD antibody interaction with S1 protein. To wrap up, in the results for ELISA applied with Ace2-Fc-coated wells, recognition of scpep8 had the potential to show slightly lower neutralization activity because of affecting the recognition of S1 protein by anti-S RBD antibody. scpep9, on the other hand, maintained the anti-S RBD antibody recognition. The effect of scpep3 on anti-S RBD antibody recognition was barely negligible since it showed mild HRP activity by giving an average OD_450_ value.

With the comparison of all results, the selected peptides had the capacity to bind directly to inactivated virus particles. According to this, QCM data showed that scpep3, scpep8, and scpep10 demonstrated the ability to bind directly to S protein. More specifically, scpep8 and scpep10 displayed proper ΔF plots that showed a decrease for each peptide addition and mass accumulation onto S protein-coated gold chip. On the other hand, scpep5 showed an unusual ΔF profile, resulting in an overall increase in ΔF, although each peptide addition decreases ΔF. In the case of scpep9, peptide addition resulted in a sharp increase in ΔF during peptide addition and a decrease in ΔF during washing steps as it removed an unbound mass of peptide molecules. For scpep12, a gradual increase in ΔF was observed in each peptide addition. To evaluate them more deeply in terms of neutralization activity, S1 proteins, which include RBD, were used as a target in ELISA. Based on the result, scpep3, scpep8, scpep8, scpep9, and scpep10 showed a neutralization activity against wild-type S1 protein. For S1 proteins of B.1.617.2 and B.1.1.2 variants, scpep3 and scpep9 showed activity for neutralizing S1 Protein binding capacity to Ace2-Fc protein for both variants. scpep8 and scpep10 also neutralized S1 protein of B.1.617.2 variant. The results were satisfying since all peptides were actually selected against inactivated virus particles, and the binding capability of peptides was evaluated for only S protein. Also, the differences between ELISA results of each variant were expected. Due to the fact that the mutations occurred in each variant, the binding as well as neutralization activity could be different. In particular, B.1.617.2 variant possesses seven different mutations and two deletions, whose two mutations in RBD, from wild-type S1 protein^36–43^. For B.1.1.2 variant, four different mutations and three deletions, whose one mutation is in RBD, occur compared with wild-type S1 protein^36,40–48^. Between B.1.617.2 and B.1.1.2 variants, only two matching mutations exist in S1 protein, which is not in RBD. With this information as a foundation, the mutations, especially in RBD could affect the neutralization activities of peptides. The differences in neutralization activities showed that scpep3 and scpep9 were the best options for inhibiting the interaction of RBD with Ace2 protein. The others possessed mild or no neutralization activities, which made them open for engineering to increase the neutralization capabilities.

Overall, SARS-CoV-2 as creating a devastating pandemic prove the fact that the pharmaceutical molecule developments are critical for this century for maintaining global health. Diverse approaches can enlighten the progress of therapeutic molecule discoveries. Selecting drug-candidate peptides by using phage display library is optimal and practical. The peptides can be developed directly into a drug molecule or can be used as developing other types of protein-based drugs. As it is, scpep3, scpep8, scpep9 and scpep10 can be used for drug development as our data showed their power for interacting with S1 protein. This approach can be also applicable for other types of infectious diseases for broaden the options for drug candidate molecules.

## Materials & Methods

### Materials

For peptide selection, Ph.D.™-12 Phage Display Peptide Library Kit (New England BioLabs□ Inc., MA, USA) was used. For protein expression, Expi293 Expression System (Gibco™) was preferred. For the interaction characterization by QCM, QSX 301 gold chips were purchased from Biolin Scientific. For HRP conjugation to S1 protein, HRP Conjugation Kit-Lightning-Link□ (Cat.#. ab102890, Abcam) was used. S1 proteins of wild-type, B.1.617.2 and B.1.1.2 were purchased from Acro Biosystems (Acro Biosystems, Cat.#SIN-C52H3 for wild-type S1; Cat.#SIN-C52Hu for B.1.617.2; Cat.#SIN-C52Hr for B.1.1.2). Anti-SARS-CoV-2 Spike RBD Neutralizing Antibody, Chimeric mAb, Human IgG1 (AM122) was obtained also from Acro Biosystems (Cat.#S1N-M122). Mouse Anti-Human IgG Fc Antibody (50B4A9) [HRP], mAb was preferred for ELISA (GenScript, Cat.#A01854). For western blotting, 6X-His Tag Monoclonal Antibody (HIS.H8) and Goat Anti-Mouse IgGH&L (HRP) were preferred (Thermo Scientific, Cat.#MA1-21315, Abcam, Cat.#ab6789, respectively). For HisTag purification, 5mL Cytiva HisTrap™ Excel column was used (Cat.#17371206).

### M13 Phage Panning for Peptide Selection

For the purpose of selecting interacting peptides against SARS-CoV-2, Ph.D.™-12 Phage Display Peptide Library Kit (New England BioLabs□ Inc., MA, USA) was preferred. The phage library supplied by the kit was amplified by following the recommended protocols of the manufacturer. The inactivated SARS-CoV-2 particles were used to coat a 96-well plate in 100 μL of coating buffer (0.1 M NaHCO_3_, pH 8.6). The coating of the plate surface was achieved by overnight incubation at 4°C. The next day, the plates were slapped to face face-down position on a clean paper towel to remove all coating buffer. For blocking of the wells, 200 μL of blocking buffer (0.1 M NaHCO_3_, pH 8.6 containing 5 mg/mL BSA) was added to each coated well. The blocking was achieved by incubation at 4°C for 2 hours. At the end of incubation, the blocking solution was removed by slapping the plate in a face-down position. To wash the wells, TBS-T (1X TBS + 0.1[v/v] Tween-20) was used, and washing was applied 6 times by adding wash buffer and removing it by slapping the plate down.

Subsequent to washing, 100 μL of 10^9^ phage particles in TBS-T was added to each well, and the plate was incubated at room temperature for 1 hour with gentle shaking. After phage-target incubation was finished, unbound phage particles were removed, and washing with TBS-T was applied 10 times by adding wash buffer and removing it by slapping the plate down onto a paper towel. When washing steps were completed, 100 μL of elution buffer (0.2 M Glycine-HCl, pH 2.2 elution buffer) was added to each well in order to disrupt phage-SARS-CoV-2 interaction, and the plate was incubated for 7 minutes at room temperature with gently shaking. Elution buffer containing phage particles was transferred to a microcentrifuge tube and neutralized by 150 μL of neutralization buffer (1M Tris-HCl, pH 9.1).

### Phage Amplification

To amplify the eluted phages in order to use the next round of panning, *E.coli* ER2738 cells were inoculated into fresh LB containing tetracycline and were grown overnight at 37°C with 200 rpm. The following day, overnight culture was diluted with a ratio of 1:100 into 20 mL fresh LB containing tetracycline and incubated at 37°C with 200 rpm until OD_600_ reached 0.01-0.05. When OD_600_ reached the desired values, eluted phage particles were added to the culture. The culture containing eluted phage particles was incubated at 37°C with 200 rpm for 4.5 hours in order to propagate phage particles obtained from panning. At the end of the incubation, the culture was centrifuged at 12000 rpm at 4°C for 10 minutes to separate cells and phage particles, which were suspended in the supernatant. The supernatant was then transferred into a new tube. For the purpose of phage precipitation, 5 mL of 20% PEG-8000/ 2.5 M NaCl solution was added to the supernatant and incubated at 4°C overnight. Subsequent to overnight incubation, the sample was centrifuged at 12000 rpm at 4°C for 15 minutes. The supernatant was removed, and finger-like phage precipitate was resuspended with 1 mL 1X TBS. For reprecipitation of phage particles, 200 μL 20% PEG-8000/ 2.5 M NaCl solution was added to phage samples in 1X TBS. The reprecipitation was achieved by incubation of samples on ice for 1 hour, which was followed by centrifugation at 14000 rpm at 4°C for 5 minutes. For this time, phage pellet was resuspended with 100 μL of 1X TBS. In order to decide the size of the amplified phage particles, phage tittering was applied. At the end of tittering protocol, each blue plaque represented a single clone of phages obtained after panning method. So as to amplify a single clone of the phages, the single blue plaque was selected with a tip and inoculated to 1 mL of *E.coli* ER2738 culture, which reached OD_600_ value between 0.01-0.05 from a dilution of overnight grown culture. The amplification of a single plaque was achieved by incubation at 37°C for 4.5 hours at 200 rpm. Next, to remove the cell pellet, centrifugation was done at 14000 rpm for 30 seconds. After the centrifugation, 800 μL of supernatant was taken from the upper part of the centrifuged samples while keeping cells undisturbed. Further, 20 μL from 800 μL single phage plaque suspension was amplified by following the same protocol applied for eluted phage amplification. All phage samples were preserved at 20°C with 50% glycerol (1:1 V/V) for long-term storage.

### Phage titering

*E.coli* ER2738 was inoculated into LB containing tetracycline antibiotic for growing cells overnight. The next day, the overnight culture was diluted with 1:100 ratio into fresh LB containing tetracycline. When OD_600_ value reached about 0.5, 200 μL of cell culture were aliquoted into sterile tubes. During this, eluted and amplified phage particles were diluted with the dilution factor ranges between 10^-2^ to 10^-4^ and 10^-8^ to 10^-12^, respectively. Each 10 μL sample from dilutions was transferred to 200 μL aliquoted cell cultures. For phage infection to occur, 1-5 minutes of incubation was applied at room temperature. At the end of the phage infection, cell-phage mixtures were added to 45°C, 3 mL top agar. Top agar containing cell-phage mixture was mixed with vortex for a couple of seconds with low rpm. Then, top agar was spread onto onto LB agar plates containing X-Gal, IPTG, and tetracycline. For observing blue/white screening, the plates were incubated at 37°C overnight. The following day, the observed blue plaques were counted in order to calculate ratio and enrichment ratio values.

### Cloning Ace2-Fc Protein and Spike Protein Encoding Gene Fragments

For the construction of AP2u2-CMV-Ace2-IgG1 plasmid and AP2u2-CMV-S _protein plasmid, AP2u2-mCherry plasmid, which was a gift from Christien Merrifield (Addgene plasmid # 27672; http://n2t.net/addgene:27672; RRID:Addgene_27672), was linearized by EcoRV and BamHI restriction enzymes^35^. With the aim of cloning IgG1, the coding sequence was amplified by PCR from pVITRO1-Trastuzumab-IgG1/κ which was a gift from Andrew Beavil (Addgene plasmid # 61883; http://n2t.net/addgene:61883; RRID:Addgene_61883)^49^. During this, cut sites of XhoI and BamHI restriction enzymes were added 5’ and 3’ ends, respectively. Human Ace2 ectodomain nucleotide sequence was chemically synthesized with the signal peptide sequence at 5’ end. In order to add EcoRV and XhoI restriction enzyme cut sites at 5’ and 3’ ends, respectively, PCR was done with appropriate Gibson assembly primers.

For S protein gene cloning, gene was chemically synthesized with EcoRV and BamHI restriction enzyme cut sites at 5’ and 3’ ends. Also, for the Gibson assembly, overhang regions were added to the ends of the gene fragment.

For the cloning of AP2u2-CMV-Ace2-IgG1, the linearized backbone, amplified IgG1, and human Ace2 ectodomain were ligated by Gibson assembly method. Also, for the cloning of AP2u2-CMV-S _protein plasmid, the linearized backbone and S protein fragment gene were ligated by Gibson assembly method. Following Gibson assembly method, the reaction product was transformed chemically into the competent strain of *E. coliDH5*α PRO by applying heat shock. After chemical transformation by applying heat shock was completed, cells were spread onto an LB agar plate containing appropriate antibiotics for the selection of colonies having cloned plasmid. The LB agar plate was incubated at 37°C overnight. The next day, grown colonies were selected, inoculated into fresh LB media containing antibiotics. The grown cultures were used for plasmid isolation, which was achieved by using GeneJET Plasmid Miniprep Kit (Thermo Fisher Scientific, Cat.#. K0502). Plasmid isolation protocol was applied according to the manufacturer’s recommendations. The isolated plasmid sequences were verified by Next Generation Sequencing (NGS) method (Intergen). After verification was completed, sequence-verified colonies were inoculated into 5 mL of LB medium with appropriate antibiotics and were grown at 37°C, 200 rpm overnight. From the overnight-grown culture, glycerol stocks were prepared with the addition of 50% glycerol into 500 μL cell culture (1:1 [v/v]). For the transfection protocol, the cells having the desired plasmids were inoculated into 50 mL LB medium containing antibiotics. The culture then, inoculated at 37°C overnight with 200 rpm shaking. The following day, cell cultures were harvested at 8000 g for 5 minutes in order to precipitate cells. After centrifugation, supernatant was removed and cell pellets were used for plasmid isolation by using NucleoBond□ Xtra Midi Plasmid DNA Purification Kit (Macherey-Nagel) according to the protocol supplied by the kit.

### Expression of Ace2-Fc and S Protein in Expi293 Expression System

Expi293 Expression System (Gibco™) was used for the expression of Ace2-Fc and S-protein. Culturing, maintenance, and transfection method of Expi293 cells were done according to the manufacturer’s recommended protocols. Basically, cell maintenance was done by growing cells at a 37°C incubator with ≥80% relative humidity and 8% CO_2_ on an orbital shaker platform at 125 rpm shake speed. For passaging, 300,000-400,000 cells/ml and 400,000-500,000 cells/ml were inoculated for 4-days incubation and 3-days incubation till the next passage, respectively. Before every passaging, the cell culture size was determined by counting cells with trypan blue with 1:1 dilution ratio. For the transfection, ExpiFectamine transfection kit was used (Gibco™). On the day prior to transfection, cells were seeded with a final density of 2.5–3 × 10^6^ viable cells/mL, and incubated overnight until transfection. On the day of transfection, the culture was diluted to a final density of 3 × 10^6^ viable cells/mL into fresh Expi Media. Meanwhile, ExpiFectamine reagent and plasmid DNA were diluted into OPTIMEM (Gibco™) according to the amount determined by the manufacturer’s transfection protocol, depending on culture volume. The diluted ExpiFectamine reagent was incubated for 5 minutes at room temperature. Then, the diluted reagent and plasmid DNA were mixed. The ExpiFectamine/DNA mix was incubated for 10-20 minutes at room temperature. At the end of the incubation, the mix was added to the culture drop by drop. The culture was incubated in a 37°C incubator with ≥80% relative humidity and 8% CO_2_ on an orbital shaker platform at 125 rpm shake speed. Following a transfection period of 18-22 hours ExpiFectamine 293 Transfection Enhancer 1 and ExpiFectamine 293 Transfection Enhancer 2 were mixed in according to the manufacturer’s protocol, in accordance with culture volume and then added to the transfected cell culture. The protein expression was achieved by incubating cells in a 37°C incubator with ≥80% relative humidity and 8% CO_2_ on an orbital shaker platform at 125 rpm shake speed for 4 days after enhancer addition.

### Ace2-Fc Protein and S Protein Purification

To obtain the highest possible protein titer, we determined the protein harvest time as five days post-transfection. On the fifth day after transfection, we centrifuged cells at 8000 g for 5 minutes at 4° C. Supernatant was transferred into a clean container. For purification, pre-equilibrated 5 mL HisTrap excel column (Cytiva, US) was used. The supernatant loaded to the column and purification was performed by using ÄKTA start protein purification system (GE Healthcare Life Sciences) according to the preferred protocols provided by the purification column manual. After samples loaded to the column, washing was performed by using wash buffer (20 mM NaH_2_PO_4_, 500 mM NaCl, 20 mM imidazole pH 7.4) for 20 column volume in order to remove excessive molecules in the supernatant other than the protein sample. Next, elution of the desired proteins was obtained by 5 column volumes of elution buffer (20 mM NaH_2_PO_4_, 500 mM NaCl, 500 mM imidazole pH 7.4). The purified proteins were concentrated by using protein concentrators. For changing buffers from elution buffer to 1X PBS pH7.4 as storage buffer, HiTrap desalting column (GE Healthcare Life Sciences) was used by connecting to ÄKTA start protein purification system. The desalting was performed by the protocol supplied with the desalting column. Pierce™ BCA Protein Assay Kit (Thermo Fisher Scientific) was used for determining the protein concentration. For long-term storage at -20°C, 10% (v/v) glycerol was added.

### SDS-PAGE and Western blotting

SDS-PAGE and western blotting were carried out to validate and characterize proteins. Protein samples were mixed with 6X SDS loading dye and boiled at 95°C for 5 minutes. Then, samples were loaded into 12% SDS-polyacrylamide gel. The electrophoresis was carried out by applying 140V for about 90 minutes. At the end of the electrophoresis, SDS-polyacrylamide gel was soaked into Coomassie blue staining solution and heated for 30 seconds in the microwave, and then shaken for 5 minutes at room temperature. In order to remove excessive dye and visualize only the protein bands clearly, the gel was transferred into destaining solution (60% ddH_2_O, 30% methanol, and 10% acetic acid). For western blot analysis, SDS-PAGE was applied with the same steps. At the end of the electrophoresis, protein samples on the gel were transferred to the activated PVDF membrane using Trans-Blot Turbo (Bio-Rad). Once the transfer was completed, the membrane was blocked for 2 hours at room temperature with 5% milk in TBS-T with gentle shaking. The blocking was followed by the next incubation step with 5% milk in TBS-T containing 1:10,000 primary anti-His mouse antibodies at 4°C overnight. At the end of the primary antibody solution incubation, the membrane was washed three times with 1X TBS-T. Proceeding from washing, the membrane was incubated with secondary antibody solution, which was prepared by 5% milk in TBS-T containing 1:10,000 horseradish peroxidase (HRP)-conjugated goat anti-mouse secondary antibodies (Abcam ab6789-1 MG). The secondary antibody incubation was applied for 1 hour at room temperature. When this step was completed, the membrane was washed three times with 1X TBS-T for the removal of excessive antibody solution. The membrane was incubated with ECL substrates (Bio-Rad 170-5060) for 1 minute at dark. Visualization of the membrane was accomplished by Vilber Fusion Solo S.

### Solid-state peptide synthesis

Rink Amide Resin (151.1 mg, 0.05 mmol) (Substitution: 0.331 meq/g) was weighed in the reactor used for peptide synthesis. The resin was washed with N, N-dimethylformamide (DMF), and then swollen by 10 mL of DMF for 20-30 minutes. After which, 5 mL of 20% piperidine was added to the reactor and waited for 3 minutes. At the end of this time interval, piperidine was removed and added again with the same volume and concentration. The second piperidine addition was waited for 10 minutes. The following step was washing, which was achieved by DMF. When the washing part is completed, Fmoc-protected amino acid (5.5 eq., 0.275 mmol) and HBTU (5.0 eq., 0.250 mmol) were measured in a test tube. 2.0 mL 0.3 M diisopropylethylamine (DIEA) in DMF was added onto them. The final mixture was placed in the reaction vessel, and the coupling mixture was processed for 1 hour. After 1 hour, the resin was washed thoroughly with DMF. This step was repeated to the point of obtaining the desired peptide elongation. When peptide elongation was completed and the last Fmoc deprotection was achieved, the resin was washed with first DMF, and then DCM, and the resin was left for drying by opened pump for 30 minutes. Next, 5 mL of cleavage cocktail (95% TFA, 5% Milli Q, 5% TIPS) was added to the reactor and incubated for 2 hours. When incubation was completed, the cleaved peptide solution was precipitated by ice-cold diethyl ether. The centrifugation was applied and precipitated peptides were washed three times with cold ether in order to remove residual small organic impurities.

### QCM Measurements for Determining Peptide-Protein Interaction

Biolin Scientific Qsense QSX301 Gold chip was preferred for QCM measurements. In order to remove possible dust or previously produced self-assembled monolayers, the gold-chip surface was cleaned by immersing it in piranha solution (H_2_O_2_/H_2_SO_4_ in 1:3 [v/v]). For the surface coating, the chip was incubated in 20 mM 110mercaptoundecanoic acid (11-MUA) at room temperature overnight. The activation of the surface was performed by EDC/NHS coupling reaction, and then protein immobilization was achieved, as previously described^50^. Each analysis was done by immobilization of 50 μg of S protein, which was followed by surface deactivation. Once the deactivation was done, 1000 μM of peptide solution was loaded into the QCM chamber for 7 minutes, and 10 minutes long washing was done afterward. The peptide addition and washing steps were repeated several times successively. During each peptide addition and washing step, frequency changes were recorded and analyzed for the first, third, fifth, seventh, and ninth overtones. The mass accumulation for each peptide onto S protein-coated chip was calculated by the formula of Δm=-C(ΔF/n), where Δm represents the mass change (ng.cm^-2^), C stands for the mass sensitivity constant, which is contingent upon the chip specification (17.7 ng.cm^-2^ for QSX 301 gold chip), ΔF depicts the change in the resonance frequency (Hz), and n signifies the number of harmonics. The mass accumulation values were evaluated depending on the values of the fifth overtones for each measurement.

### Neutralizing Activity Determination with ELISA

The synthesized peptide candidates were analyzed by ELISA assay. Due to this, 96-well ELISA plates were coated with 100 μL of 2.5 μg/mL purified Ace2-Fc in 1X PBS, pH 7.4 at +4°C overnight. The following day, Ace2-Fc coating solution was removed, and wells were washed with 300 μL 1X PBS-T once. After washing, blocking of wells was achieved by incubating the plate with 200 μL of blocking buffer (3% [w/v] BSA, 3% [w/v] skimmed milk in 1X PBS-T) at room temperature for 1 hour. When the incubation period ended, 100 μL of stabilizing buffer (3% [w/v] mannitol, 3% [w/v] sucrose in 1X PBS-T) was added to wells and continued to incubation at room temperature for 1 hour. For activity analysis, S1 proteins of wild-type, B.1.617.2, and B.1.1.2 variants were conjugated with horseradish peroxidase (HRP) by using HRP Conjugation Kit-Lightning-Link®. Simultaneously, 30 μL of 120 ng/mL HRP-labeled S1 proteins of wild-type, B.1.617.2, and B.1.1.2 variants in stabilizer were mixed with 30 μL of 1 mM peptide solution in 1X PBS, pH 7.4. The HRP-labeled S protein-peptide mix was incubated at 37°C for 1 hour. At the end of the incubation for both plate and peptide-protein mix, 50 μL of HRP-labeled S protein-peptide mix was added to the wells, and incubation was done at room temperature for 1 hour. Once the incubation was completed, washing was accomplished by using 300 μL 1X PBS-T five times. When, 1X PBS-T was removed from wells, 100 μL TMB was added to wells, and wells were incubated at room temperature for 15 minutes at dark environment. For halting the reaction, 100 μL 0.16 M H_2_SO_4_ was added to wells. OD values were measured at 450 nm and 630 nm by using BioTek EL808 Microplate Reader, via GEN5 software.

## Acknowledgements

Conflict of Interest

## Supplementary Materials

**Supplementary Table:** The properties of peptides

**Supplementary Figure S1:** Western blotting results of Ace2-Fc and S protein

**Supplementary Figure S2:** QCM ΔF records for S protein immobilizations

**Supplementary Figure S3:** Experimental design for ELISA

## Notes

### Competing Interest Statement

The authors have declared no competing interest.

